# The megabase-sized fungal genome of *Rhizoctonia solani* assembled from nanopore reads only

**DOI:** 10.1101/084772

**Authors:** Erwin Datema, Raymond J.M. Hulzink, Lisanne Blommers, Josè Espejo Valle-Inclan, Nathalie van Orsouw, Alexander H.J. Wittenberg, Martin de Vos

## Abstract

The ability to quickly obtain accurate genome sequences of eukaryotic pathogens at low costs provides a tremendous opportunity to identify novel targets for therapeutics, develop pesticides with increased target specificity and breed for resistance in food crops. Here, we present the first report of the ~54 MB eukaryotic genome sequence of *Rhizoctonia solani,* an important pathogenic fungal species of maize, using nanopore technology. Moreover, we show that optimizing the strategy for wet-lab procedures aimed to isolate high quality and ultra-pure high molecular weight (HMW) DNA results in increased read length distribution and thereby allowing generation of the most contiguous genome assembly for *R. solani* to date. We further determined sequencing accuracy and compared the assembly to short-read technologies. With the current sequencing technology and bioinformatics tool set, we are able to deliver an eukaryotic fungal genome at low cost within a week. With further improvements of the sequencing technology and increased throughput of the PromethION sequencer we aim to generate near-finished assemblies of large and repetitive plant genomes and cost-efficiently perform *de novo* sequencing of large collections of microbial pathogens and the microbial communities that surround our crops.

## Introduction

Global food production suffers from pests and diseases causing yield losses of up to 80% (Oerke and Dehne, 2004). Currently, mankind relies heavily on the use of pesticides in agricultural settings, but their usage is increasingly restricted due to concerns about pesticide retention in food crops, effects on off-target organisms and risk of resistance in major pathogen/insect groups. In contrast to pesticides, host resistance that has a genetic basis provides a sustainable solution to pest control. The rapid and low cost generation of accurate genome sequences of pathogenic micro-organisms will allow superior target site prediction for a better understanding of the molecular *modus operandi* of pests and diseases during infection of crops. In turn, this understanding is of crucial importance in identifying resistant sources for breeding of genetic resistances in crops.

The basidiomycetous plant-pathogenic fungus *Rhizoctonia solani* Kühn is an economically damaging soil-born pathogen that infects seeds and seedlings and causes damping-off diseases in a wide range of crop species, including maize, rice, soybean, potato and sugar beet (González et al. 2006). *R. solani* is a species complex represented by distinct anastomosis groups (AGs) that can be further divided into subgroups featuring different growth characteristics, host plant ranges, and genetics. To date, high-quality genome sequence information is available for four *R. solani* isolates represented by three different AGs and isolated from different host plants (Zhang et al., 2013; Cubeta et al., 2014; Hane et al., 2014; Wibberg et al., 2015). Comparative analysis of the draft genome sequences revealed the existence of genomic differences across different isolates including genome size and gene constitution, likely resembling differences in host specificity and life cycle (Wibberg et al. 2015). However, the genome sequences of these isolates are highly fragmented due to the short-read sequence technologies used to generate the data. Long-read sequencing, such as offered by Pacific Biosciences (PacBio) and Oxford Nanopore Technologies (ONT), allows direct sequencing of DNA fragments of tens of kilobases in size without the need for clonal amplification (reviewed by Goodwin *et al.* 2016). In contrast to other sequencing technologies, the read length of nanopore sequences is therefore limited primarily by the length of the DNA molecules themselves. As such, this technology holds the potential to generate ultra-long and unbiased reads of megabases in size, enabling the resolution of long repeats and facilitating the accurate assembly of complex genomic regions. Despite the inherent ability of nanopore sequencing for generating *de novo* assemblies of large, complex genomes, the use of nanopore reads in recent literature has been limited to *de novo* assembly of small-sized, prokaryotic genomes (Deschamps et al. 2016, Karamitros et al. 2016, Loman et al, 2015, Szalay and Golovchenko, 2015) and medium-sized eukaryotic genomes following a hybrid assembly approach (Goodwin et al. 2015).

Recently, a preliminary report of *de novo* genome assemblies of multiple yeast isolates generated from nanopore reads was published (Istace et al, 2016). Here, we have assembled a complex fungal genome from nanopore reads only. Upon isolation of ultra-pure high molecular weight (HMW) DNA from *R. solani*, we were able to produce a highly contiguous draft genome sequence from a moderate sequencing depth. Compared to previous assemblies, we have reduced the number of contigs by an order of magnitude and increased the N50 contig size to just below 200 kb. Our results indicate that high-quality, nearly finished eukaryotic genomes can be achieved with moderate efforts and at low cost.

## RESULTS

To facilitate *R. solani* genome assembly using nanopore reads only, we aimed at maximizing the read length derived from the MinION. Thereto, we used size-selection to exclude small DNA fragments from the samples and thus extracted genomic DNA of high integrity. This resulted in HMW DNA from 20 Kb in length and with a mass middle over 50 Kb (Figure S1). The DNA was further purified to facilitate precise and reproducible shearing of the DNA prior to library preparation. To determine the effect of read length on assembly contiguity, three long-insert nanopore libraries that differ in their mean fragment length were synthesized (Figure S2). Two libraries were prepared from randomly sheared genomic DNA with a mean fragment length of ~13 kb and ~19 kb and one library was synthesized from intact genomic DNA. All DNA samples and corresponding libraries showed a discrete and narrow fragment size distribution without the presence of small DNA fragments. The three libraries were sequenced using seven R7.3 flow cells producing a total of 77,799 2D pass reads that together correspond to 834 megabases with an average read length of 10.7 kb. We observed a strong positive effect of library fragment size on read length distribution, with the majority of long reads produced by the non-sheared library (Figure S2). The reads were *de novo* assembled, using canu, into 606 contigs spanning 54 Mb with an N50 contig length of 199 kb, thereby providing the most contiguous *R. solani* assembly to date and the largest eukaryotic genome generated from nanopore reads only to date (Table 1).

**Table 1:** Genome assembly features of sequenced *R. solani* isolates.

| Description | AG1-IA | AG1-IB | AG3 | AG8 | AG5 |
| --- | --- | --- | --- | --- | --- |
| Reference | Zheng *et al.*, 2013 | Wibberg *et al.*, 2015 | Cubeta *et al.*, 2014 | Hane *et al.*, 2014 |  |
| Sequencing technologies | Illumina GAII | 454 GS-FLX + Illumina MiSeq | Sanger + 454 GS-FLX | Illumina HiSeq | ONT MinION |
| Number of contigs | 6,452 | 3,793 | 6,040 | 7,606 | 606 |
| N50 contig length (kb) | 20 | 35 | 26 | 7 | 199 |
| Assembled genome size (Mb) | 36.9 | 42.8 | 51.7 | 39.8 | 54 |
| Mitochondrial genome size (kb) | 147 | 163 | 236 | 140 | 254 |

In order to verify the quality of the nanopore-derived *R. solani* genome assembly, we compared the sequence with an assembly obtained from a single paired-end MiSeq run. A total of 13.9 million merged read pairs with an average fragment length of 360 nt were obtained from a single MiSeq run and assembled into 123,016 contigs with total length 71 Mb and an N50 contig length of 1,029 nt. Subsequent comparison of the MiSeq assembly to the nanopore assembly identified 99,147 single nucleotide differences, 68,500 insertions and 146,383 deletions relative to the nanopore assembly in 48 Mb of aligned sequence. Alignment of the individual MiSeq reads to the nanopore assembly followed by variant calling resulted in 24,697 single nucleotide differences, 71,829 insertions and 181,551 deletions over the entire 54 Mb of the nanopore assembly. Assuming the MiSeq data to be of perfect quality, this provides a lower bound on the error rate of the nanopore assembly of one substitution error per 2,186 bases, one insertion error per 700 bases and one deletion error per 297 bases. The higher number of single nucleotide differences observed from the whole-genome alignment results from the greater length of the MiSeq assembly, which implies that multiple MiSeq contigs can be aligned to a single position on the nanopore assembly.

The histogram of 31-mer frequency derived from the MiSeq reads (Figure S3) displays two major peaks representing the heterozygous and homozygous fractions of the genome, respectively. From these data, the haploid genome size of *R. solani* was estimated to be 46.6 Mb with 38.6 Mb of unique sequence and 8 Mb of repetitive sequence per haploid genome. Strikingly, our nanopore assembly spans 54 Mb and displays a bimodal read depth distribution when the MiSeq reads are mapped against it, with many individual contigs corresponding to one of the two read depth peaks (Figure S4 and S5). The first peak corresponds to approximately 22 Mb of heterozygous sequence (*i.e.* 11 Mb of haploid sequence that is assembled in two copies) and the second peak corresponds to approximately 29 Mb of homozygous sequence (*i.e.* 29 MB of haploid sequence that is indistinguishable between the two copies). This suggests that that the homozygous fraction of the genome is primarily assembled in a single copy, whereas the heterozygous fraction is present as two separate copies.

To determine the contribution of ultra-long reads to the contiguity of the assembled contigs, three additional *de novo* assemblies were generated from the nanopore data using the ultra-fast miniasm assembler: one assembly starting from all reads; one assembly with 75% of total data volume represented by the shortest reads (“miniasm-short”); and one assembly with 75% of total data volume represented by the longest reads (“miniasm-long”). The resulting assemblies are summarized in Table 2. Compared to Canu, miniasm produces a smaller but more contiguous assembly. Whereas the total assembled genome differs minimally between the two assemblies derived from 75% of the data volume, the number of contigs nearly doubles (and accordingly, the N50 contig length nearly halves) when using shorter reads, compared to longer reads. Despite the overall inferior results of the miniasm assemblies, these data clearly demonstrate the impact of ultra-long DNA molecules on assembly contiguity.

**Table 2:** Comparison of nanopore assembly statistics.

| Description | canu | miniasm | miniasm-short | miniasm-long |
| --- | --- | --- | --- | --- |
| Number of contigs | 606 | 313 | 505 | 322 |
| Assembled genome size (Mb) | 54.0 | 44.7 | 37.9 | 39.8 |
| Average contig length (kb) | 89 | 143 | 75 | 124 |
| Median contig length (kb) | 39 | 67 | 46 | 73 |
| N50 contig length (kb) | 199 | 294 | 125 | 216 |
| Maximum contig length (kb) | 1,673 | 1,616 | 718 | 1,168 |

As is evident from Table 1, the data presented here represents the most contiguous assembly to date. Comprehensive genome annotation is outside the scope of the current study; nonetheless, 85% (10,736 out of 12,616) genes from the AG1-IB isolate could be located on the here-produced AG5 assembly. This includes 2,428 out of 2,520 “core genes” described by Wibberg et al (2015). Moreover, *R. solani* is known to harbour a strikingly large and complex mitochondrial genome compared to other filamentous fungi (Losada et al., 2014). Our assembly recovered the mitochondrial genome in a single contig of 254,189 nt without any effort targeted towards obtaining this result, highlighting once more the strength of ultra-long DNA reads for genome reconstruction. Both the nuclear and mitochondrial genome sequences of the AG5 isolate display a high level of sequence divergence compared to those of previously sequenced isolates (Figure S6). This corresponds well to the high level of sequence variation that was demonstrated between both the nuclear and mitochondrial genomes of those isolates (Wibberg et al, 2015; Losada et al, 2014).

## DISCUSSION

The results described in this paper demonstrate the strength of HMW DNA purification combined with long-read nanopore sequencing to produce a highly contiguous genome sequence of a complex eukaryotic plant pathogen with moderate efforts and at low cost. Even at this early stage of nanopore sequencing technology we have succeeded in mapping the complex genome of an economically important plant pathogen, which could not be readily performed with short-read technologies: the N50 contig size of our assembly is up to 28 times larger when compared to assemblies from short-read data only, and up to 6 times larger when considering genomes assembled using multiple sequencing platforms (Table 1). Such superior genomes assemblies will aid – amongst others – in improved identification of genes, which is hampered or obscured in genomes of lesser quality. Whereas the error rate of our current assembly is higher than preferred, we expect future improvements to nanopore sequencing technology and the chemistry used therein to resolve these issues. In fact, novel R9 technology has significantly increased sequence output and accuracy (Instace et al, 2015). Given the promising results of the here-described approach, we aim to use our ability to generate ultra-long sequence reads from HMW DNA fragments, and leverage the increased throughput of the PromethION sequencer to generate near-finished assemblies of large and repetitive plant genomes in the near future. The continuous advancements in nanopore sequencing technology, sequencing throughput, accuracy as well as bioinformatics applications to analyze the type of data will enable cost-efficient *de novo* sequencing of large collections of microbial pathogens, the complex genomes of the crops they infect and the microbial communities that they inhabit with the aim to increase food safety and breed for disease resistance in the world’s staple crops.

## MATERIALS AND METHODS

### Fungal culturing and tissue collection

*Rhizoctonia solani* AG5 (strain number CBS 339.84) isolated from *Zea mays* was obtained from CBS-KNAW Fungal Biodiversity Centre (Utrecht, the Netherlands) and cultured on potato dextrose agar (PDA; Sigma-Aldrich, Zwijndrecht, the Netherlands) plates at 8°C. For DNA extraction, 0.5 cm^2^ agar blocks colonized with *R. solani* were used to inoculate 200 mL liquid potato dextrose medium. The cultures were incubated at 24°C and 130 rpm for four to five days. Mycelial mats were collected and washed twice with demineralized water followed by two washes with 500 mM NaCl_2_ / 50 mM EDTA pH 8.0. After washing, the mycelia was collected and the excess of mositure was squeezed out. The dried tissue was collected and stored at −80°C for later use.

### DNA extraction and purification

Deep-frozen mycelium tissue was ground into a fine powder in liquid nitrogen. The powder was resuspended in four volumes of DNA extraction buffer (100 mM tris-HCl pH8, 70 mM EDTA pH8.0, 2% (v/v) SLS, 2% (v/v) 2-mercaptoethanol, 100 μg/mL proteinase K) and the homogenate was incubated at 55°C for 1 h. After incubation, the homogenate was centrifuged for 10 min at 5,087 *g* and the supernatant was extracted with 1 volume of chloroform / isoamylalcohol (24:1). DNA was precipitated with isopropanol and the dried DNA pellet was resuspended in 100 μL of demineralized water. DNA quantity and purity was determined with the Nanodrop UV-Vis spectrophotometer (Thermo Scientific, Wilmington, NC, USA) and the Qubit 2.0 fluorometer (Thermo Fisher Scientific, Waltham, MA, USA). Integrity assessment of the genomic DNA was performed using an 2200 TapeStation Instrument (Agilent Technologies, Santa Clara, CA, USA) along with Genomic DNA ScreenTapes. To obtain ultra-pure HMW DNA, the DNA was further purified and concentrated with the Nucleic Acid Extraction System of Boreal Genomics (Boreal Genomics, Vancouver, BC Canada) following the manufacturer's instructions.

### DNA size selection and size determination

To remove short fragment DNA, purified genomic DNA was size selected using the BluePippin preparation system (Sage Science, Beverly, MA, USA) following the instructions in the user guide for high-pass DNA size selection. Purified DNA was sized-selected with the 0.75% DF 50 kb Marker S1 High-Pass 6-10 Kb protocol choosing a BP start and end cutoff of 12 and 65 Kb, respectively (target size of 38.5 Kb). After size selection, the sample was concentrated with 1x volume Agencourt AMPure beads (Beckman Coulter Inc., Brea, CA, USA) following the manufacturer's instructions. The effect of the sizing was assessed by electrophoretic analysis with the 2200 TapeStation instrument (Agilent Technologies, Santa Clara, CA, USA) and by pulse-field gel electrophoresis (PFGE). For size determination on the PF gel, 1.5 μg PFGE standard λ ladder DNA (Bio-Rad Laboratories Inc., Hercules, CA, USA) and 1 μg of CHEF DNA Standards DNA (Bio-Rad Laboratories Inc., Hercules, CA, USA) were loaded. Electrophoresis was performed in 0.5x TBE buffer with a CHEF-DRII system (Bio-Rad Laboratories Inc., Hercules, CA, USA) using the following electrophoresis conditions: linear switch time ramp, two state mode, initial switch time of 1 s, final switch time of 12 s, run time of 9.5 h, reorientation angle of 120°, voltage gradient of 12 V/cm, and run temperature of 14°C.

### Nanopore library preparation and MinION sequencing

For two libraries, about 1.5 μg of HMW genomic DNA was converted into 12.5 Kb and 18.8 Kb fragments through hydrodynamic shearing using a MegaRuptor^®^ DNA shearing system (Diagenode) following the manufacturer's instructions. After shearing, the length of the DNA fragments was determined using an 2200 TapeStation Instrument (Agilent Technologies, Santa Clara, CA, USA) along with Genomic DNA ScreenTapes. Prior to library preparation, the sheared DNA was concentrated with 1x volume of Agencourt Ampure XP beads (Beckman Coulter Inc., Brea, CA, USA). Preparation of the ONT libraries was performed using the Genomic DNA Sequencing kit version SQK-MAP006 according to the instructions of Oxford Nanopore Technologies (Oxford, UK). In contrast to the protocol, no control DNA (CS DNA) was added during the library preparation. Both the size distribution and concentration of the final libraries (named presequencing library) was determined with the 2200 TapeStation Instrument (Agilent Technologies, Santa Clara, CA, USA) using a Genomic DNA ScreenTape. For the 17.9 Kb presequencing library, three R7.3 flow cells were loaded containing respectively 53.5 ng, 53.5 ng, and 89.2 ng library DNA. For both the 21.3 and 56.6 Kb libraries, two R7.3 flow cells were used containing 37.8 ng and 125.2 ng (21.3 kb) and 7.8 ng and 28.6 ng (56.6 kb) library DNA. All runs were performed on the MinION MK1 device for 48 h using the 48h_sequencing_run_FLO_MAP103.py protocol (MinKNOW version 0.51.1.62 b201602101407). The basecalling was performed in the cloud using the 2D Basecalling for SQK-MAP006 protocol (rev. 1.69; Metrichor version v.2.38.10).

### MiSeq library preparation and sequencing

From 1 μg of genomic DNA Illumina fragment libraries were prepared with an average insert size of 350 bp according to the TruSeq^®^ DNA PCR-Free Library Prep protocol (Part # 15036187 Rev. D). DNA was sequenced (250 bp Paired End) on a MiSeq using V3 reagents. Supplementary Table 1 details the number of reads produced.

### Sequence assembly and analysis

Nanopore sequence data was extracted from the FAST5 files output by Metrichor^TM^ using poretools (Loman and Quinlan, 2014) and subsequently assembled with miniasm (Li, 2015) and Canu (Berlin et al, 2015). Post-assembly error correction was performed using Nanopolish (Loman et al, 2015). MiSeq sequence data was processed using FLASH (Magoc and Salzberg, 2011) to merge overlapping read pairs from short fragments; non-overlapping read pairs were discarded. Merged MiSeq reads were aligned to the Canu assembly using BWA MEM (Li, 2013). Read alignments were further processed with Picard (http://broadinstitute.github.io/picard/) and samtools (Li, 2011) after which variants were called using GATK Lite (De Pristo et al, 2011). Variants with read depth below 20 or above 200 were discarded from subsequent analyses. *De novo* assembly of merged MiSeq reads was performed with SOAPdenovo2 (Luo et al, 2012). The *R. solani* genome size was estimated using GenomeScope (<u>https://github.com/schatzlab/genomescope</u>) from k-mer counts obtained through Jellyfish2 (Marcais & Kingford, 2011). Assembled genome sequences were compared using dnadiff from the MUMmer suite (Kurtz et al, 2004). Gene sequences from AG1-IB were aligned to the AG5 assembly with BLASTN from the BLAST° suite of programs (Camacho et al, 2008). Alignment covering less than 50% of the gene sequence or having a sequence identity lower than 50% were discarded.

## Supplemental Tables and Figures

**Figure S1:**
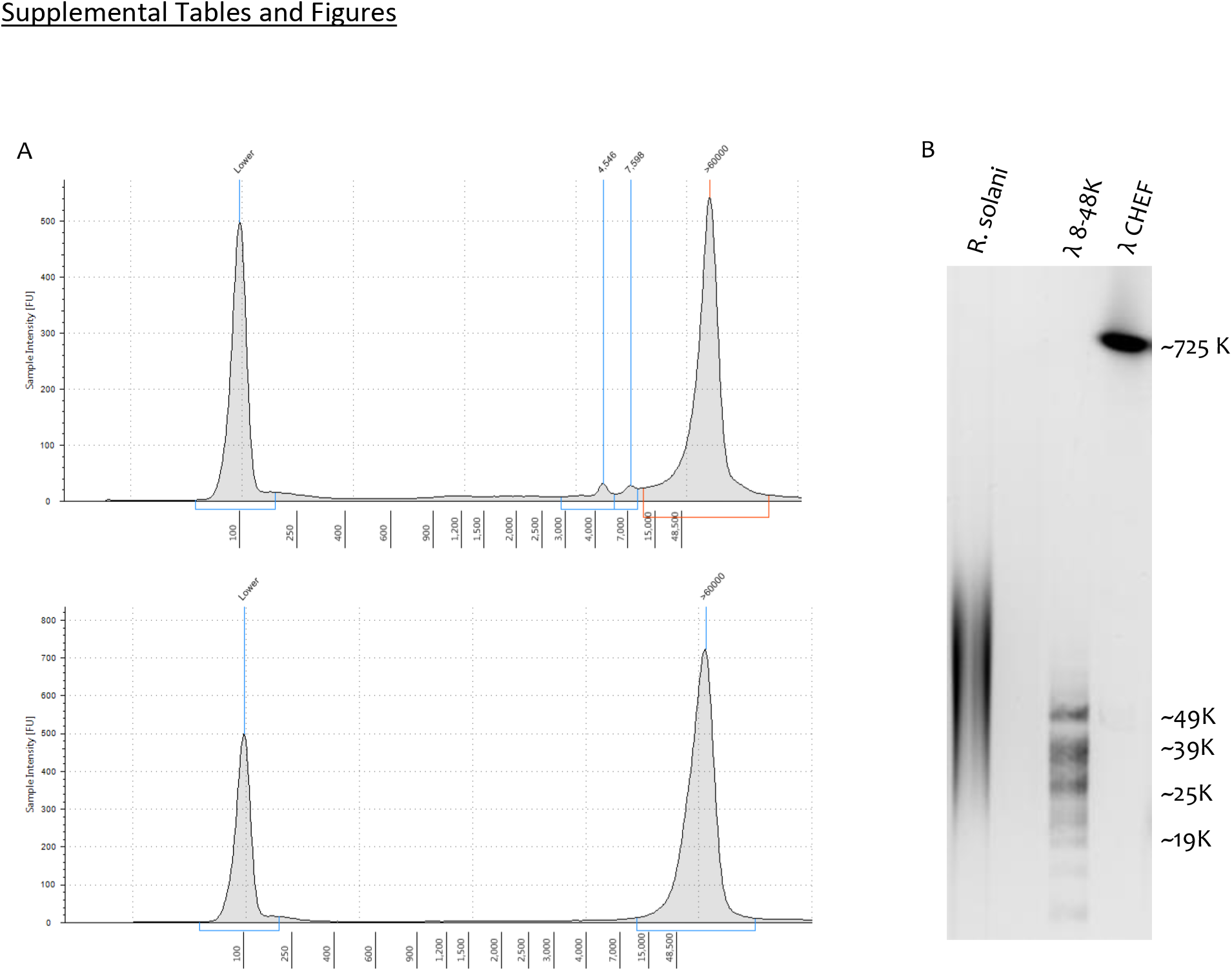
The effect of size selection on the size distribution of *R. solani* genomic DNA. Electropherograms derived from the Agilent Tapestation showing the size distribution of genomic DNA before (upper graph) and after (lower graph) size selection (A). The Y axis shows the signal intensity (FU) and the X-as shows the DNA fragment size in basepairs. The 100 bp peak in the electropherograms is derived from the size marker. Although non-sized genomic DNA showed a discrete and narrow sized distribution from 60 Kb, a low level of short fragment DNA could be observed. Size selection resulted in the complete removal of the low molecular weight DNA and the recovery of the high molecular weight DNA. Pulse-field gel electrophoresis revealed that most of the sized high molecular weight DNA was over 20 Kb in length and had a mass middle of about 50 – 60 Kb (B).

**Figure S2:**
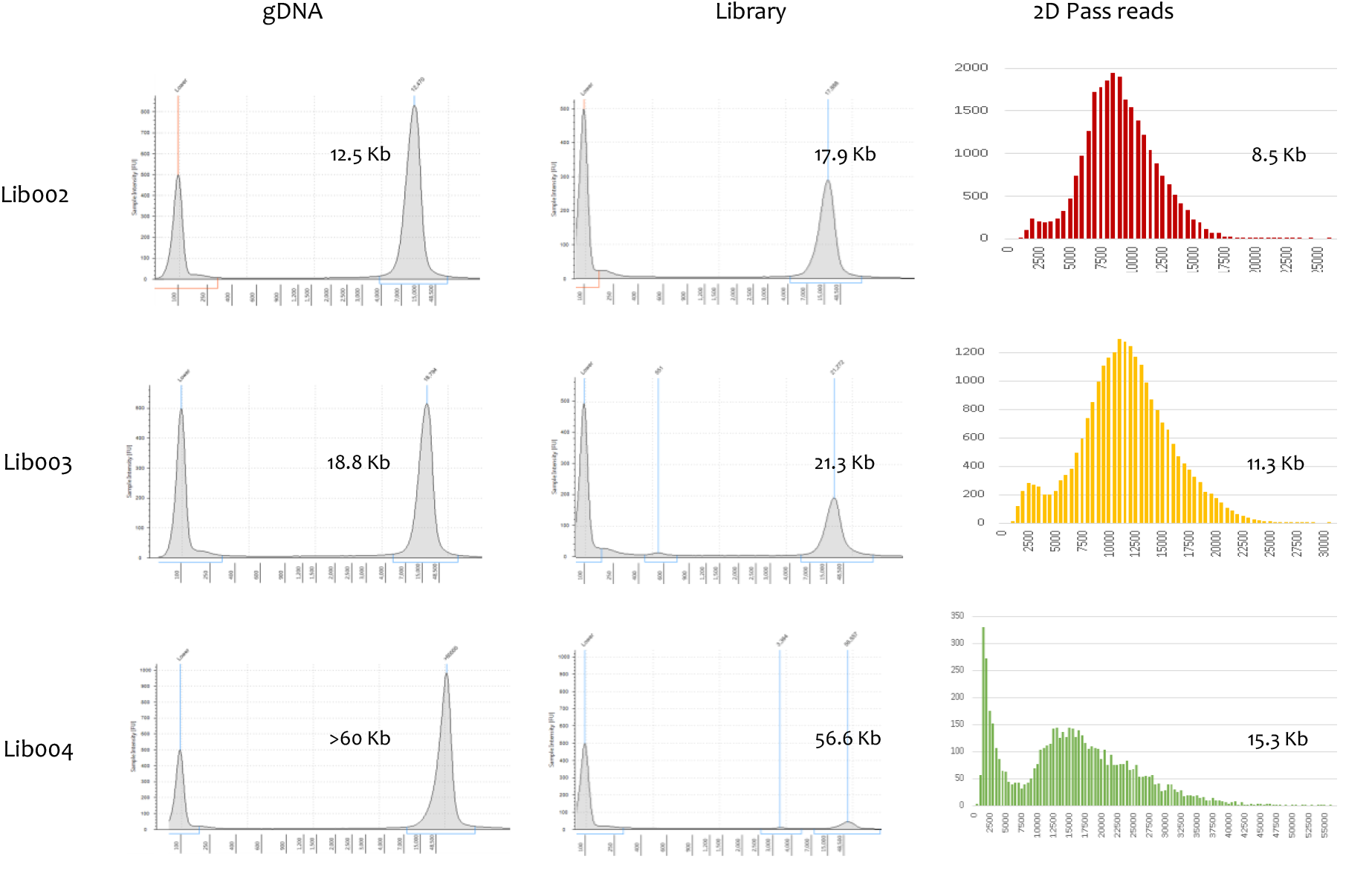
Size distribution of sheared *R. solani* DNA and corresponding nanopore libraries and 2D pass reads. Electropherograms derived from the Agilent Tapestation showing the size distribution of sheared and intact genomic DNA (graphs at the left) and the corresponding nanopore libraries (graphs in the middle). The signal intensity (FU) and DNA fragment size in basepairs are depicted at the Y axis and X axis, respectively. The 100 bp peak in the electropherograms is derived from the size marker. The graphs at the right show the size distribution of the corresponding 2D reads. Although the size distribution between the size-selected genomic DNA samples and the corresponding libraries are comparable, a substantial difference in size distribution could be observed between the DNA samples / nanopore libraries and the corresponding 2D pass reads.

**Figure S3:**
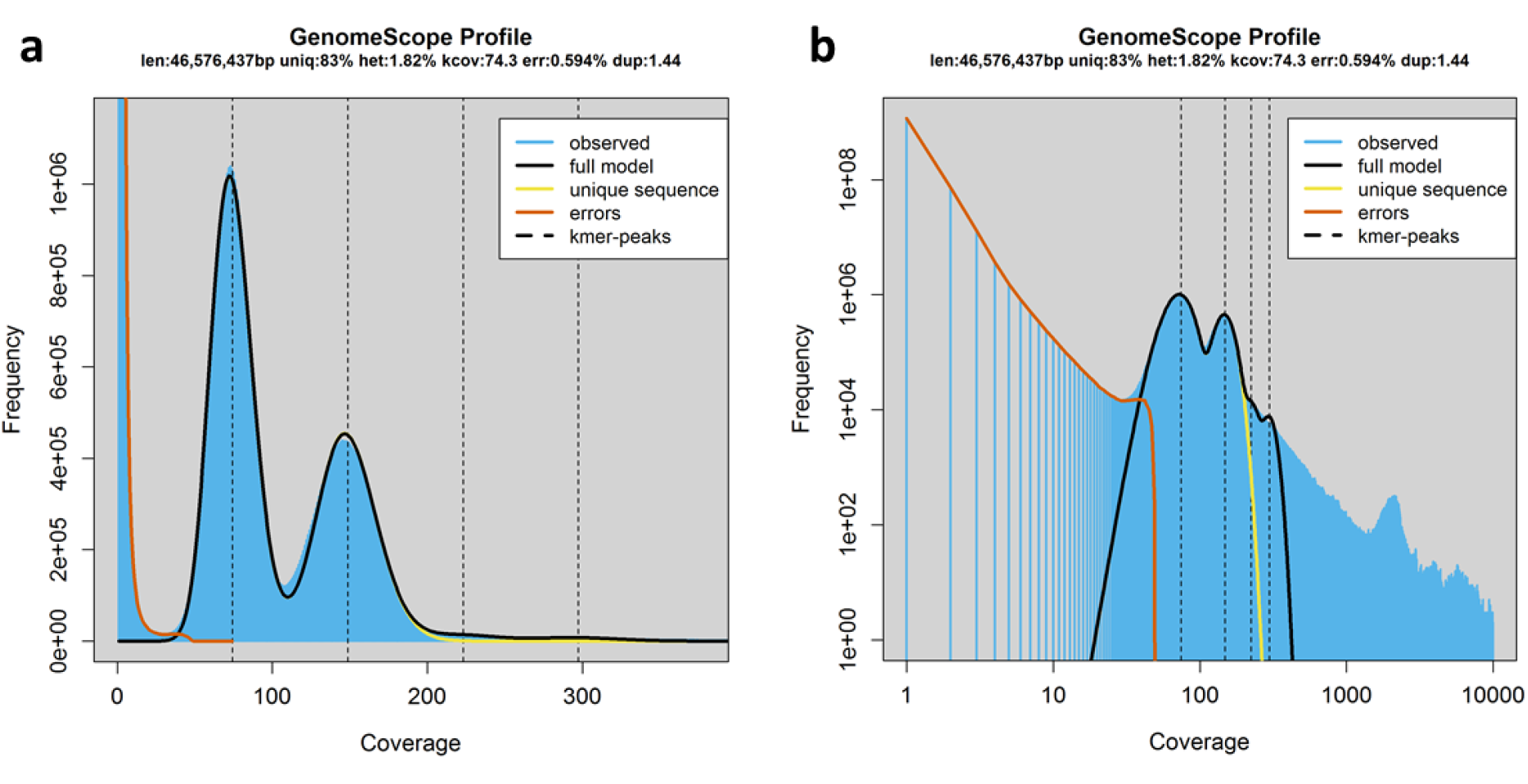
Abundance of 31-mers in MiSeq data as determined by GenoScope in (**a**) linear and (**b**) log_10_ scale. The x-axis represents 31-mer counts; the y-axis shows the number of 31-mers in the MiSeq read data with that abundance.

**Figure S4:**
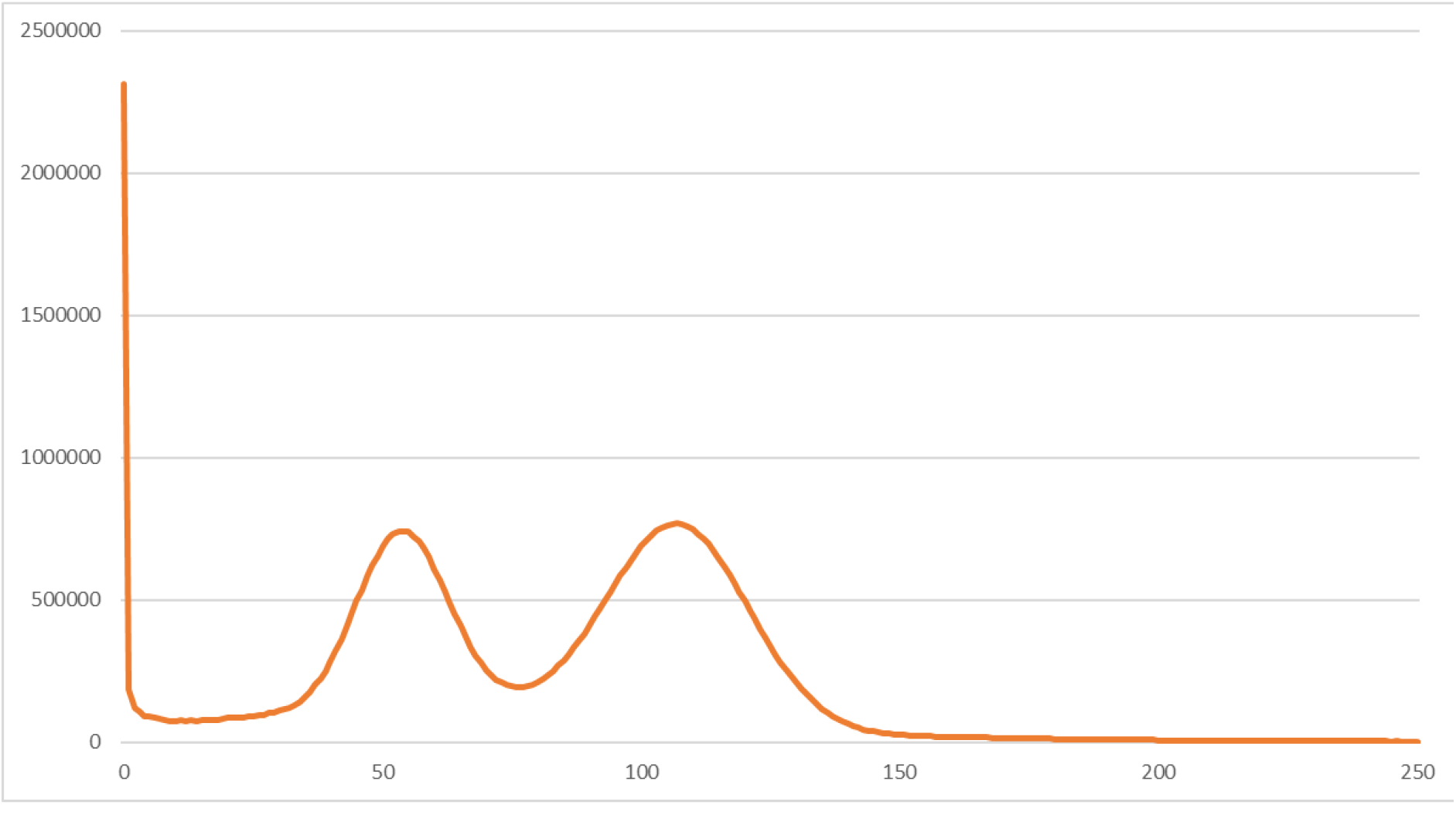
Read depth distribution of MiSeq reads mapped against the Canu assembly of the nanopore reads, after polishing. The x-axis represents read depth; the y-axis shows the number of genome positions with that number of reads mapped against it. In total, 2.3 Mb out of the 54 Mb assembly have no MiSeq reads mapped against it. The heterozygous and homozygous read depth peaks are at 54x and 107x, respectively.

**Figure S5:**
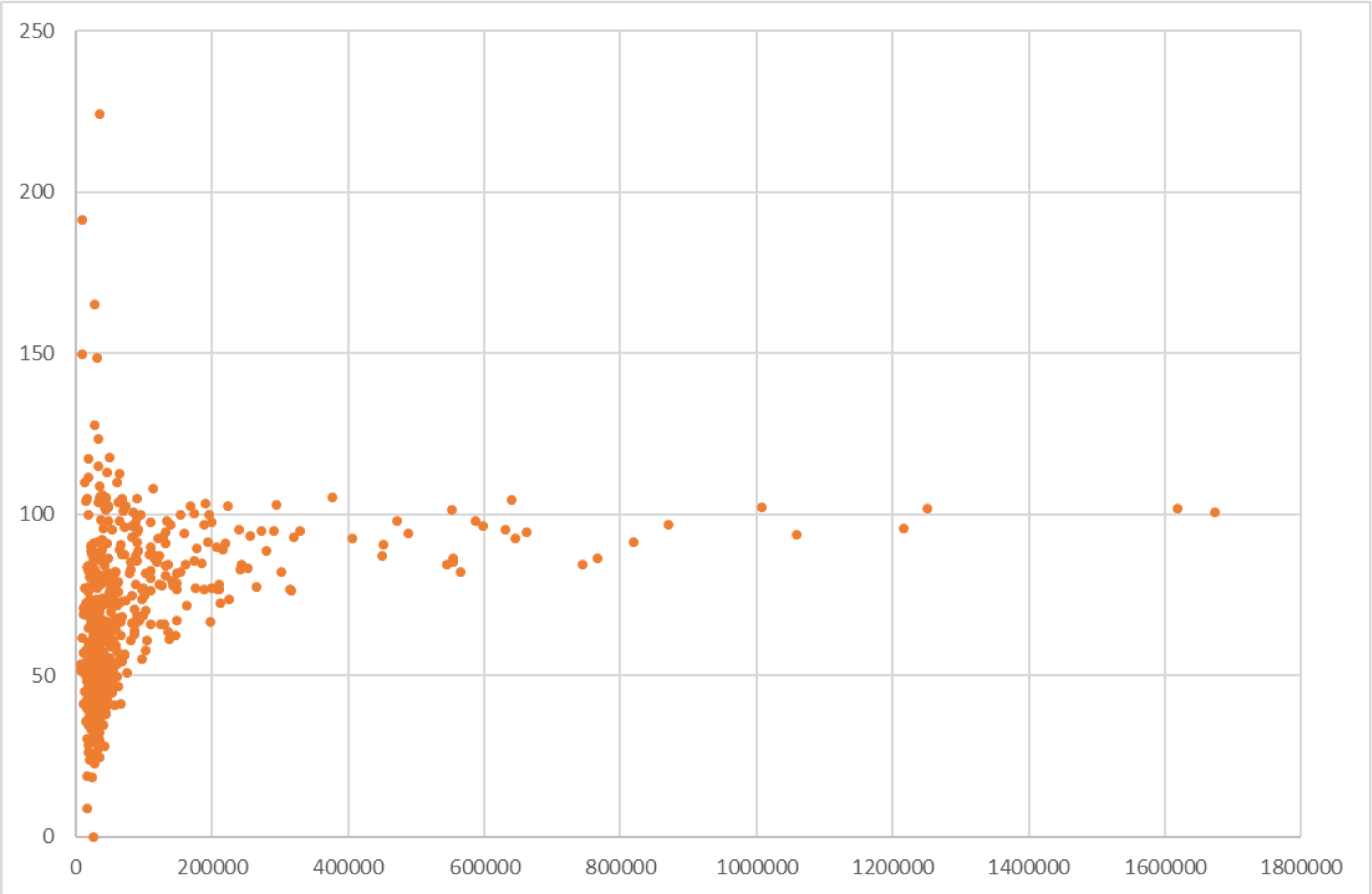
Average read depth of MiSeq reads (y-axis) versus contig length (x-axis) in the Canu assembly of the nanopore reads. Each data point represents a contig,. The majority of contigs larger than 500 kb have a read depth corresponding to the second peak in Supplementary Figure EDA2 and represent the homozygous fraction of the genome, whereas many small contigs have approximately half that read depth and represent the heterozygous fraction. The y-axis has been truncated at 250; a single contig with an average read depth of 1,306x is not shown.

**Figure S6:**
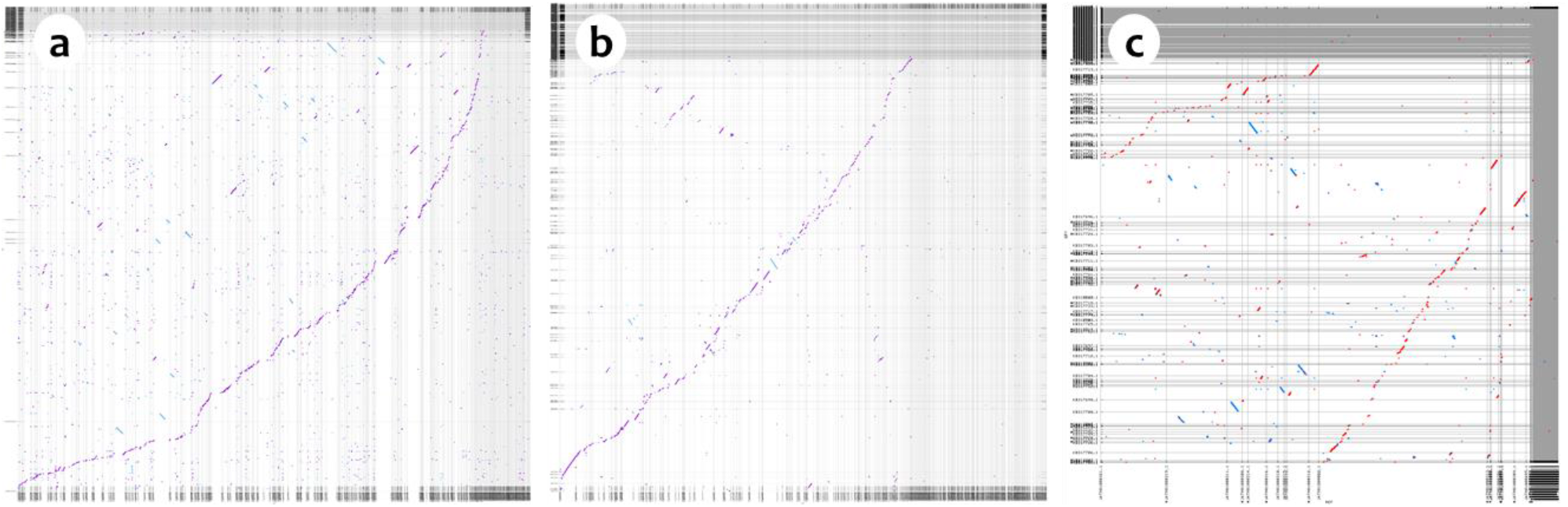
Whole genome alignments between different isolates of *R. solani*. (**a**) AG3 (Cubeta et al, 2014; y-axis) versus the Canu assembly of AG5 (x-axis); (**b**) AG1-IA (Zheng et al, 2013; y-axis) versus the Canu assembly of AG5 (x-axis); (**c**) AG1-IA (y-axis) versus AG3 (x-axis). Red lines represent direct matches; blue line represent matches between opposite strands. Dotplots generated with mummer-plot (Kurtz et al, 2004).

## Literature

Berlin K., Koren S., Chin C.S., Drake P.J., Landolin J.M., and Phillippy A.M. (2015) Assembling Large Genomes with Single-Molecule Sequencing and Locality Sensitive Hashing. Nature Biotechnology; 33(6):623–30

Camacho C., Coulouris G., Avagyan V., Ma N., Papadopoulos J., Bealer K., and Madden T.L. (2008) BLAST+: architecture and applications. BMC Bioinformatics 10:421.

DePristo M., Banks E., Poplin R., Garimella K., Maguire J., Hartl C., Philippakis A., del Angel G., Rivas M.A., Hanna M., McKenna A., Fennell T., Kernytsky A., Sivachenko A., Cibulskis K., Gabriel S., Altshuler D., and Daly M., (2011) A framework for variation discovery and genotyping using next-generation DNA sequencing data. NATURE GENETICS 43:491–498

Deschamps S., Mudge J., Cameron C., Ramaraj T., Anand A., Fengler K., Hayes K., Llaca V., Jones T.J., and May G. (2016) Characterization, correction and de novo assembly of an Oxford Nanopore genomic dataset from Agrobacterium tumefaciens. Scientific Reports 6, 28625

González García V., Portal Onco M. A., and Rubio Susan V. (2006) Review. Biology and Systematics of the form genus Rhizoctonia. Spanish Journal of Agricultural Research 4(1), 55–79

Goodwin S., Gurtowski J., Ethe-Sayers S., Deshpande P., Schatz M.C., and McCombie, W.R (2015) Oxford Nanopore sequencing, hybrid error correction, and de novo assembly of a eukaryotic genome. Genome Research 25(11): 1–7

Goodwin S., McPherson J.D., and McCombie W.R. (2016) Coming of age: ten years of next-generation sequencing technologies. Nature Reviews Genetics 17, 333–351

Istace B., Friedrich A., d'Agata L., Faye S., Payen E., Beluche O., Caradec C., Davidas S., Cruaud C., Liti G., Lemainque A., Engelen S., Wincker P., Schacherer J., and Aury J.-M. (2016). De novo assembly and population genomic survey of natural yeast isolates with the Oxford Nanopore MinION sequencer. BioRxiv 066613

Karamitros T., Harrison I., Piorkowska R., Katzourakis A., Magiorkinis G., and Lutamyo Mbisa J., (2016) De Novo Assembly of Human Herpes Virus Type 1 (HHV-1) Genome, Mining of Non-Canonical Structures and Detection of Novel Drug-Resistance Mutations Using Short- and Long-Read Next Generation Sequencing Technologies. PLOS One 11(6), e0157600.

Karlsson E., Lärkeryd A., Sjödin A., Forsman M., and Stenberg, P. (2015) Scaffolding of a bacterial genome using MinION nanopore sequencing. Scientific Reports 5, 11996

Kurtz S., Phillippy A., Delcher A.L., Smoot M., Shumway M., Antonescu C., and Salzberg S.L. (2004) Versatile and open software for comparing large genomes. Genome Biology 5:R12

Li H., (2011) A statistical framework for SNP calling, mutation discovery, association mapping and population genetical parameter estimation from sequencing data. Bioinformatics 27(21):2987–93.]

Li H., (2013) Aligning sequence reads, clone sequences and assembly contigs with BWA-ME. http://arxiv.org/abs/1303.3997

Li H., (2015) Minimap and miniasm: fast mapping and de novo assembly for noisy long sequences. http://arxiv.org/abs/1512.01801

Loman N.J., and Quinlan A.R., (2014) Poretools: a toolkit for analyzing nanopore sequence data. Bioinformatics 30(23):3399–401

Loman N.J., Quick J., Simpson J.T. (2015) A complete bacterial genome assembled de novo using only nanopore sequencing data. Nat Methods 12(8):733–735.

Luo R., Liu, B., Xie Y., Li Z., Huang W., Yuan J., He G., Chen Y., Pan Q., Liu Y., Tang J., Wu G., Zhang H., Shi Y., Liu Y., Yu C., Wang B., Lu Y., Han C., Cheung D.W., Yiu S.-M., Peng S., Xiaoqian Z., Liu G., Liao X., Li Y., Yang H., Wang J., Lam T.-W., and Wang J., (2012) SOAPdenovo2: an empirically improved memory-efficient short-read de novo assembler. GigaScience 2012 1:18.

Magoc T., and Salzberg S.L., (2011) FLASH: fast length adjustment of short reads to improve genome assemblies. Bioinformatics. 2011 Nov 1; 27(21): 2957–2963.

Marçais G., and Kingsford C., (2011) A fast, lock-free approach for efficient parallel counting of occurrences of k-mers. Bioinformatics 27: 764–70.

Oerke E.C., and Dehne H.W. (2004) Safeguarding production - losses in major crops and the role of crop protection. Crop Protection 23: 275–285.

Szalay T., and Golovchenko J.A. (2015) De novo sequencing and variant calling with nanopores using PoreSeq. Nature Biotechnology 33, 1087–1091

